# Mechanistic models of signaling pathways deconvolute the functional landscape of glioblastoma at single cell resolution

**DOI:** 10.1101/858811

**Authors:** Matías M. Falco, María Peña-Chilet, Carlos Loucera, Marta R. Hidalgo, Joaquín Dopazo

**Affiliations:** Clinical Bioinformatics Area, Fundación Progreso y Salud (FPS), Hospital Virgen del Rocío, Sevilla, 41013, Spain; Bioinformatics in Rare Diseases (BiER), Centro de Investigaciones Biomédicas en Red en Enfermedades Raras (CIBERER).; Unidad de Bioinformática y Bioestadística, Centro de Investigación Príncipe Felipe (CIPF).; Functional Genomics Node, FPS/ELIXIR-ES, Hospital Virgen del Rocío, Sevilla, Spain

## Abstract

The rapid development of single cell RNA-sequencing (scRNA-seq) technologies is revealing an unexpectedly large degree of heterogeneity in gene expression levels across the different cells that compose the same tissue sample. However, little is known on the functional consequences of this heterogeneity and the contribution of individual cell-fate decisions to the collective behavior of the tissues these cells are part of. Mechanistic models of signaling pathways have already proven to be useful tools for understanding relevant aspects of cell functionality. Here we propose to use this mechanistic modeling strategy to deconvolute the complexity of the functional behavior of a tissue by dissecting it into the individual functional landscapes of its component cells by using a single-cell RNA-seq experiment of glioblastoma cells. This mechanistic modeling analysis revealed a high degree of heterogeneity at the scale of signaling circuits, suggesting the existence of a complex functional landscape at single cell level. Different clusters of neoplastic glioblastoma cells have been characterized according to their differences in signaling circuit activity profiles, which only partly overlap with the conventional glioblastoma subtype classification. The activity of signaling circuits that trigger cell functionalities which can easily be assimilated to cancer hallmarks reveals different functional strategies with different degrees of aggressiveness followed by any of the clusters.

In addition, mechanistic modeling allows simulating the effect of interventions on the components of the signaling circuits, such as drug inhibitions. Thus, effects of drug inhibitions at single cell level can be dissected, revealing for the first time the mechanisms that individual cells use to avoid the effect of a targeted therapy which explain why and how a small proportion of cells display, in fact, different degrees of resistance to the treatment. The results presented here strongly suggest that mechanistic modeling at single cell level not only allows uncovering the molecular mechanisms of the tumor progression but also can predict the success of a treatment and can contribute to a better definition of therapeutic targets in the future.

## Introduction

Since the beginning of the century, transcriptomic technologies, which evolved from microarrays (1) to RNA-seq (2), have provided and increasingly accurate insight into mRNA expression (3). The technological advances of RNA-seq technologies have increased the resolution in the quantification of transcripts until the unprecedented level of the mRNA component of individual single cells. The possibility of studying gene expression at the single cell level opens the door to novel biological questions which were not possible using current tissue-level RNA sequencing approaches. For example, single-cell RNA-sequencing (scRNA-seq) has allowed a high-resolution analysis of developmental trajectories (4,5), the detailed characterization of tissues(6,7), the identification of rare cell types (8), or the analysis of stochastic gene expression and transcriptional kinetics (9,10), just to cite a few cases.

The continuous publication of scRNA-seq studies is producing an increasingly large wealth of data on cell-level gene activity measurements under countless conditions. However, the functional consequences of such gene activity at single cell level remains mostly unknown. Among the many methods and applications published for the management of scRNA-seq data (11), only a small proportion of them provide some functional insights on the results. For example, MetaNeighbor (12), SCDE (13) or PAGODA (14), annotate cell types based on conventional gene set enrichment analysis (15,16). Other algorithms, such as SCENIC (17), PIDC (18), SCODE (19) or SINCERITIES (20), offer the possibility of inferring regulatory networks as well. However, functional profiling methods have evolved from the analysis of simple gene sets or inferred regulatory gene networks to more sophisticated computational systems biology approaches that allow a mechanistic understanding on how molecular cell signaling networks enables cells to make cell-fate decisions that ultimately define a healthy tissue or organ, and how deregulation of this signaling network lead to pathologic conditions (21–23). Specifically, mechanistic models have helped to understand the disease mechanisms behind different cancers (24–27), rare diseases (28,29), the mechanisms of action of drugs (29,30), and other physiologically interesting scenarios such as obesity (31) or the post-mortem cell behavior of a tissue (32). Although there are several mechanistic modeling algorithms available that model different aspects of signaling pathway activity, *Hipathia* (24) has demonstrated to outperform other competing algorithms in terms of sensitivity and specificity (23).

Here, we propose the use of mechanistic models of signaling activities (24,33) that trigger cell functionalities related with cancer hallmarks (34), as well as other cancer-related relevant cellular functions to understand the consequences of gene expression profiles on cell functionality at single cell level. Such mechanistic models use gene expression data to produce an estimation of activity profiles of signaling circuits defined within pathways (24,33). An additional advantage of mechanistic models is that they can be used not only to understand molecular mechanisms of disease or of drug action but also to predict the potential consequences of gene perturbations over the circuit activity in a given condition (35). Actually, in a recent work, our group has successfully predicted therapeutic targets in cancer cell lines with a precision over 60% (25).

An interesting model to be studied from the viewpoint of mechanistic models is glioblastoma, the most common and aggressive of gliomas (36). Glioblastoma current treatment includes maximal safe surgical resection followed by radiotherapy and chemotherapy (37), often combined with other drugs such as bevacizumab in an attempt of overcoming resistances (38). Despite this intense treatment, the mean survival of glioblastoma patients is only 15 months and resistances to the therapy are quite common (39–41). This high rate of treatment failure has been attributed to the lack of specific therapies for individual tumor types (42,43). Moreover, it is well known that glioblastoma tumors with a common morphologic diagnosis display a high heterogeneity at the genomic level (44).

The availability of glioblastoma single cell gene expression data (45) provides a unique opportunity to understand the behavior of a cancer type at the cell level. Here, we show for the first time how mechanistic models applied at single-cell level provide an unprecedentedly detailed dissection of the tumor into functional profiles at the scale of individual cells that throw new light on how cells ultimately determine its behavior. Moreover, since mechanistic models allow simulating interventions on the system studied, we show a comprehensive simulation of the potential effect of drugs at single cell level that disclose, for the first time, the mechanisms and strategies used by subpopulations of cells to evade the effect of the drug.

## Results

### Selection of the optimal imputation method

Since mechanistic models take into account the topology of signaling circuits to estimate signal transduction activity in the cell, the discrimination between genes with missing expression values and genes that are not expressed is crucial, given that, depending on the location of the gene within the circuit, it can play the role of a switch. Since dropout events (the observation of a gene at a moderate expression level in one cell that cannot be detected in another cell) are quite common in scRNA-seq experiments (46), and taking them as zero values can cause dramatic effects on the inferred activity of the circuit, the use of imputation methods is crucial for the application of the mechanistic model. Among the best performer imputation methods available (47), three of them were checked to decide which one is optimal in the context of signaling pathway activity inference: MAGIC (48), SCimpute (49) and Drimpute(50).

In order to decide what imputation method produced the most realistic results we used the clustering produced in the original single cell glioblastoma study (45) as ground truth. There, the authors applied tSNE (51) over the 500 most variable and highly expressed genes and then cluster the resulting data with k-means. They found 12 main clusters with a homogeneous cell composition that was further experimentally validated, which were: Astrocytes, two Myeloid cell clusters, three Neoplastic cell clusters, Neurons, Oligodendrocytes, OPCs and three Vascular cell clusters. Then, gene expression values were imputed using the above-mentioned methods (MAGIC, SCimpute and Drimpute). Next, gene expression values were used to infer signaling circuit activities with the *Hipathia* algorithm (24) as implemented in the Bioconductor application (52). The values of circuit activity were subjected to the same procedure (tSNE dimensionality reduction and k-means clustering) and the resulting clusters were compared to the original ones obtained in the glioblastoma study using the Rand index (53). Figure 1A shows the clustering obtained with the genes following the procedure described above (equivalent to the Figure 2 of the original study (45)), which can be compared with the clustering of the samples using the circuit activities obtained with the gene expression values imputed with SCimpute (Figure 1B), Drimpute (Figure 1C) and MAGIC (Figure 1D) The comparison of the clusters obtained with the three imputation methods were SCImpute: 0.745, DrImpute: 0.852 and MAGIC: 0.858. Although MAGIC rendered a slightly better rand coefficient, Drimpute was chosen as imputation method because the dispersion of the clusters obtained was very similar to the one observed in the ground truth clustering (Figure 1A).

**Figure 1.**
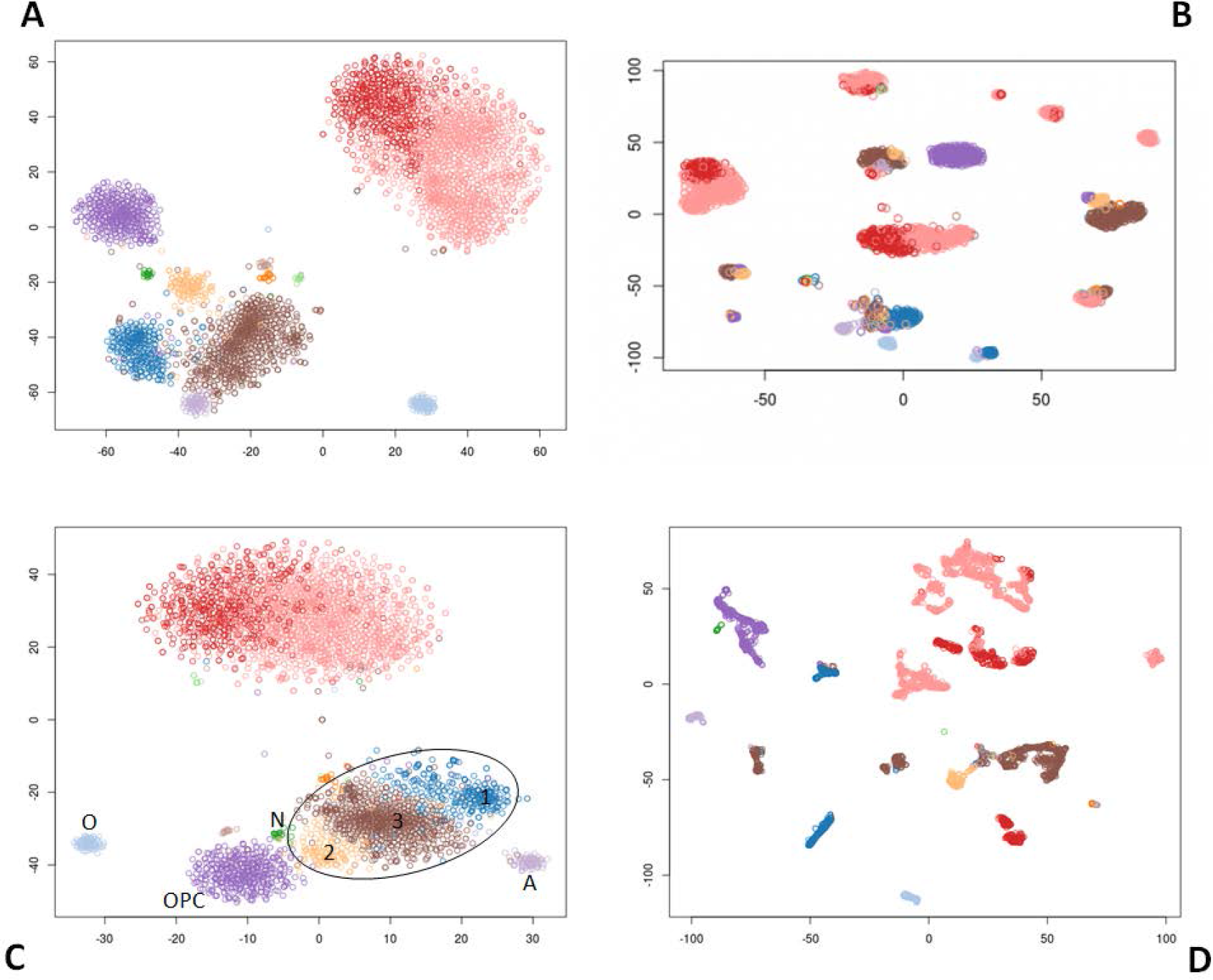
Clustering of the samples based on gene expression and on signaling circuit activities obtained with different gene imputation methods. A) shows the clustering obtained with the gene expression values following the procedure described in the original glioblastoma study (44), Clustering obtained using all the circuit activities inferred using gene expression values imputed with B) SCimpute, C) DrImpute and D) and MAGIC. In panel C, label O indicates the oligodendrocyte cluster, label OPC indicates the oligodendrocyte precursor cell cluster, label A indicates the astrocyte cluster and label N marks a small cluster of neurons. The ellipse approximately delimits the neoplastic cells among which three sub-clusters are distinguished, labeled with consecutive numbers. Unlabeled cells are a mixture of vascular and myeloid cells.

**Figure 2.**
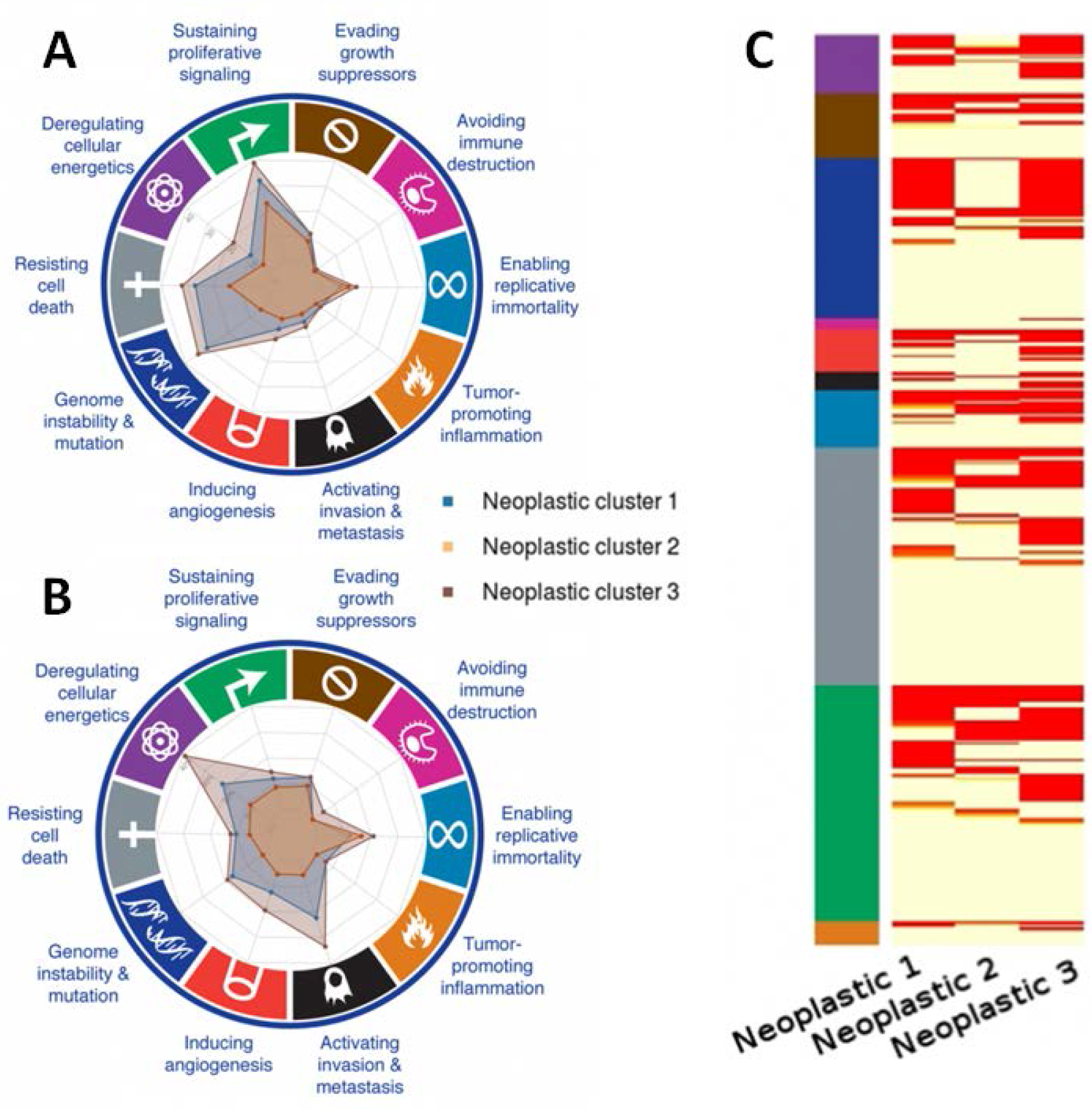
Circuits related to cancer hallmarks observed in the three neoplastic cell clusters. A) absolute number of circuits with a significant differential activity in the neoplastic cells; B) percentage of circuits with a differential activity with respect to the total number of circuits annotated to any of the cancer hallmarks; C) Heatmap with the signaling circuits related to the different cancer hallmarks that have been found to be differentially activated in cells of each neopalstic cluster.

### Functional characterization of cancer cells

Once verified that cell types defined by gene expression profiles (45) are supported by signaling profiles as well, the obvious comparison is the glioblastoma cell clusters versus the clusters composed by the different brain cells (oligodendrocytes, neurons, astrocytes and OPCs). Since circuits activity bridges gene expression to signaling activity and ultimately cell functionality, an assessment of the differences between cell types from a functional perspective can be achieved by means of a differential cell activity statistical contrast. The cell functional responses triggered by the circuits differentially activated can be easily retrieved, and among them, those related with cancer hallmarks (34) can be identified using the *CHAT* tool (54), as explained in methods.

In order to detect which circuits display a significant change in activity, the three neoplastic cell clusters (1, 2 and 3 in Figure 1C) are compared to the normal cells brain cells (oligodendrocytes, oligodendrocyte precursor cells, astrocytes and neurons, labeled as O, OPC, A and N, respectively, in Figure 1C).

The comparison between the neoplastic clusters and the brain normal cells resulted in two different patterns of circuit activity: neoplastic clusters 1 and 3 present a higher number of signaling circuits differentially activated (309 and 336, respectively) than neoplastic cluster 2 (only 96 circuits). Figure 2 represents the number of differentially activated signaling circuits involved in cancer hallmarks observed in the three neoplastic cell clusters. This representation provides a summary of the strategy used by any particular neoplastic cluster in terms of the number of signaling circuits that control cell functionalities identifiable as cancer hallmarks. Figure 2A depicts the absolute number of circuits with a significant differential activity in the neoplastic cells and Figure 2B the same results but as percentages with respect to the total number of circuits annotated to any of the cancer hallmarks. Table 1 summarizes the number of signaling circuits related to cancer hallmarks common to the three clusters (first column) and specific for each cancer type (subsequent columns). The common functional signature of this cancer is clearly driven by circuits related to *Resisting cell death*, *Sustaining proliferative signaling* and *Enabling replicative immortality* hallmarks, completed with circuits related to *Evading growth suppressors*, *Inducing angiogenesis* and *Tumor promoting inflammation* hallmarks. From Figure 1 it becomes apparent that neoplastic clusters 1 and 3 are using a functional strategy different from that used by neoplastic cluster 2. The first two ones display a functional signature compatible with a more aggressive behavior: they have many extra circuits related to *Resisting cell death*, *Sustaining proliferative signaling* hallmarks but, in addition, both clusters have circuit activity related to *Deregulation of cellular energetic*, *Genome instability and mutation* as well as *Invasion and metastasis* hallmarks (see Table 1 and Figure 2C for a detail). Conversely, neoplastic cluster 2 does not seem to have much more extra functional activity beyond the common functional signature, which suggests a less aggressive character, especially because of the absence of circuit activity related to cell energetic or to invasion and metastasis.

**Table 1.** Summary of the different functional strategies followed by the different cells in the three neoplastic clusters in terms of the circuits differentially activated with respect to the normal tissue.

| Cancer hallmark | Common circuits | Neoplastic cluster 1 | Neoplastic cluster 3 | Neoplastic cluster 2 |
| --- | --- | --- | --- | --- |
| Resisting cell death | 4 | 13 | 14 | 2 |
| Sustaining proliferative signaling | 8 | 14 | 14 | 2 |
| Deregulating cellular energetics |  | 4 | 8 |  |
| Genome instability and mutation |  | 5 | 6 |  |
| Inducing angiogenesis | 2 | 2 | 5 |  |
| Enabling replicative immortality | 5 | 1 | 3 |  |
| Activating invasion and metastasis |  | 2 | 3 |  |
| Evading growth suppressors | 3 | 3 | 2 |  |
| Tumor promoting inflammation | 1 | 1 | 1 |  |
| Avoiding immune destruction |  |  | 1 |  |

### Functional based stratification of glioblastoma cells

Neoplastic clusters have been defined according to the individual profiles of signaling circuit activities observed for each cell. The advantage of this way of cell stratification is that the functional profiles of each group are well defined. Current glioblastoma classification stratifies tumors into three subtypes, classical, proneural, and mesenchymal, from less to more aggressive, based on the signature of 50 genes (55). The *SubtypeME* tool from the GlioVis data portal (56) was used to assign subtype to each individual cell. Interestingly, when cells of the three neoplastic clusters are typed, the distribution of markers is very coherent with their functional activity profiles. Thus, Neoplastic cluster 2 is mainly composed by cells belonging to the classical subtype (see Table 2), in coincidence with its functional profile less aggressive. On the other hand, Neoplastic cluster 2 has an important component of proneural cells, as well as a smaller proportion of mesenchymal cells, which is coherent with its more aggressive functionality triggered by its signaling activity, which includes modifications in circuits related to cell metabolism, genomic instability and metastasis. Moreover, the functional profile of Neoplastic cluster 3 seems to be even more aggressive than that of Neoplastic cluster 1. This group of glioblastoma cells not only has more circuits related to the same hallmarks that Neoplastic cluster 1 but also has circuits that trigger functionalities for Avoiding immune destruction (Table 2 and Figure 1). It is interesting noticing that, although the conventional stratification in Classical, Mesenchymal and Proneural classes is illustrative of the behavior of the cells it does not completely fit with the stratification based on whole cell functional profiles.

**Table 2.** Distribution of the different glioblastoma subtypes across the three neoplastic cell clusters.

|  | Classical | Mesenchymal | Proneural |
| --- | --- | --- | --- |
| Neoplastic cells 1 | 92 | 44 | 135 |
| Neoplastic cells 2 | 107 | 3 | 15 |
| Neoplastic cells 3 | 540 | 141 | 14 |

### Effect of a drug at single cell level

Mechanistic models can be used to simulate the effect of an intervention over the system studied (25,35). Specifically, single cell transcriptomic data offers, for the first time, the possibility of modeling the effects of a targeted drug at the level of individual cells.

The current indication for glioblastoma patients treatment is temozolomide which induces DNA damage, that can be combined with other drugs such as bevacizumab to overcome resistances (38). Moreover, bevacizumab, which is indicated for several advanced cancer types, has recently been suggested for glioblastoma targeted treatment (57–59). Actually, the effect of bevacizumab, a humanized murine monoclonal antibody targeting the vascular endothelial growth factor ligand (*VEGFA*), can easily be simulated in the mechanistic model. *VEGFA* gene participates in 6 pathways (*VEGF signaling pathway, Ras signaling pathway, Rap1 signaling pathway, HIF-1 signaling pathway, PI3K-Akt signaling pathway* and *Focal adhesion*) and is part of 81 circuits, 39 of them directly related to cancer hallmarks (18 to *Resisting cell death*, 9 to *Sustaining proliferative signaling*, 4 to *Genomic instability and mutation*, 3 to *Evading growth suppressors*, 2 to *Enabling replicative immortality*, 2 to *Inducing angiogenesis* and 1 to *Deregulation of cellular energetic*). As described in methods, the inhibition of *VEGFA* can be simulated by taking the gene expression profile of a single cell, creating a simulated profile by setting the inhibited gene to a low value and comparing two profiles (24,35).

Figure 3 shows the impact of the inhibition of *VEGFA* on the different cells in terms of changes in the activities of signaling circuits in which this protein participates. The Y axis depicts the magnitude of this change in the activities of signaling circuits. There are clearly two different behaviors in the response: most of the cells present a drastic change in many signaling circuit activities (responders) while a smaller number of them present a much lower affectation on them (low-responders). Interestingly, most of the cells with a low-response to bevacizumab are typed as Mesenchimal.

**Figure 3.**
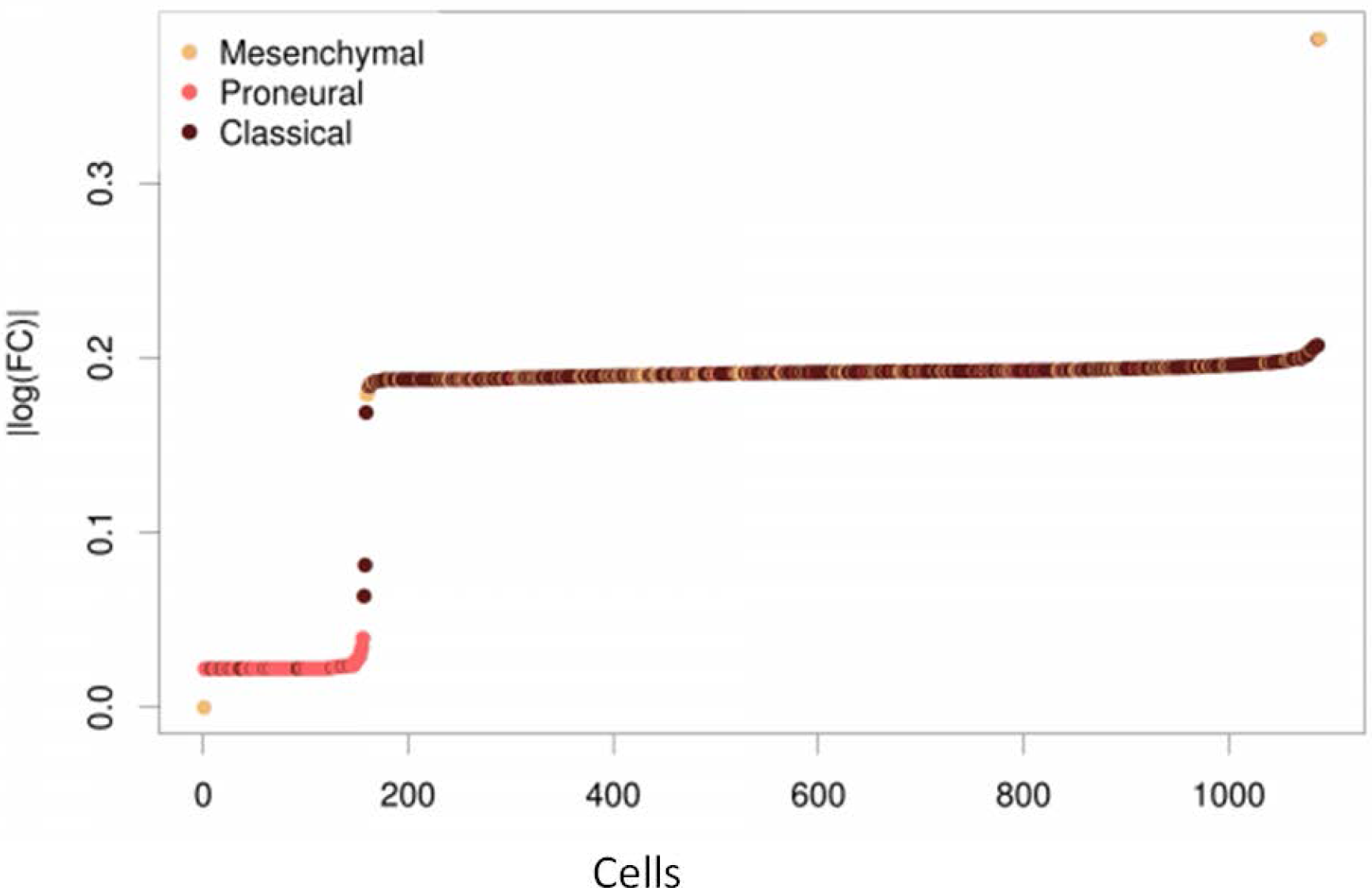
Impact of the inhibition of *VEGFA* by bevacizumab over the different neoplastic cells in terms of changes in the activities of signaling circuits in which this protein participates. The Y axis depicts the magnitude of this change in the activities of signaling circuits.

A close look at the consequences of the inhibition of *VEGFA* in different cells provides an interesting explanation for the observed differences. *VEGFA* is upstream in the chain of signal transduction in several circuits of different pathways. In the circuits within *Ras signaling pathway, Rap1 signaling pathway* and *PI3K-Akt signaling pathway,* the *VEGFA* protein potentially share the role of signal transducer with other 40 proteins. Figure 4A clearly depicts how the balance between the expression level of *VEGFA* in the responsive cells and *KITLG* and *FGF2* proteins, that can take a similar signaling role, changes. The inhibition of *VEGFA* in the responsive cells will radically inhibit the signal. However, the low-responsive cells have already *VEGFA* at low expression levels and the signal is transmitted by *KITLG* and *FGF2* instead, which ultimately compromises the success of the drug. A similar scenario occurs with the *Focal adhesion pathway*, in which *VEGFA* shares the signal transduction role with other 12 proteins. In this case, the low-responder cells are characterized by a low level of *VEGFA* compensated with a high level of *PDGFD*, which makes these signaling circuits in the low-responding cells virtually insensitive to the inhibition of *VEGFA* (see Figure 4B). Only in the case of 6 signaling circuits belonging to the *HIF signaling pathway* and *VEGF signaling pathway* the protein *VEGFA* is the only signal transducer in the node. In this case, low-responder cells have this circuit constitutively down and, consequently, are not affected by the inhibition (Figure 4C).

**Figure 4.**
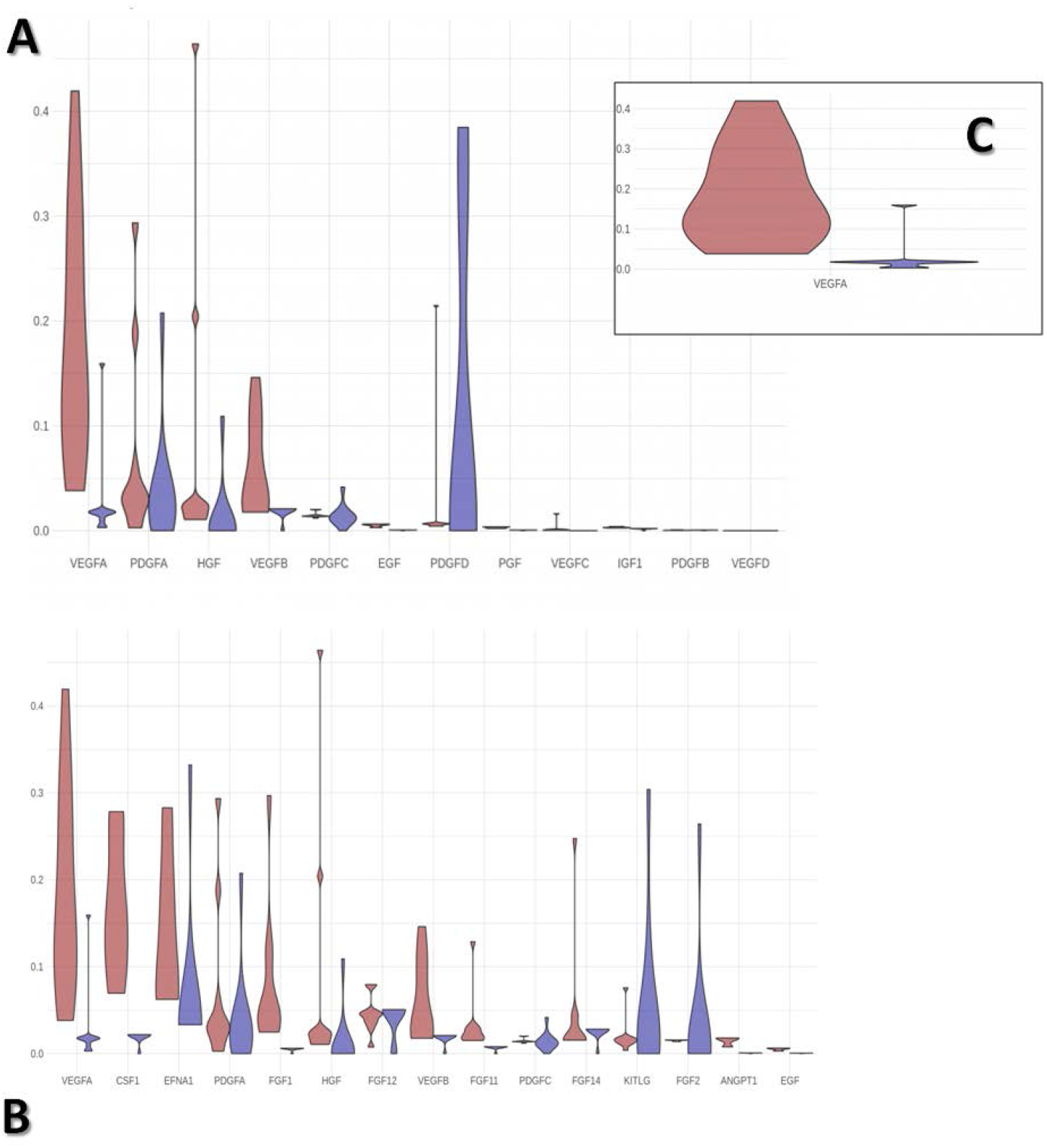
Distribution of the values of normalized gene expression in the genes located within the effector node of the different signaling circuits affected by the bevacizumab inhibition. The distribution of observed expression levels in responder cells appears in red and in the low-responders in blue. A) in the receptor node of the circuits within *Ras signaling pathway, Rap1 signaling pathway* and *PI3K-Akt signaling pathway,* the *VEGFA* protein potentially share the role of signal transducer with other 40 proteins (*CSF1*, *EFNA1*, *PDGFA*, *FGF1*, *HGF*, *FGF12*, *VEGFB*, *FGF11*, *PDGFC*, *FGF14*, *KITLG*, *FGF2*, *ANGPT1*, *EGF*, *PDGFD*, *EFNA5*, *ANGPT2*, *PGF*, *VEGFC*, *FGF18*, *EFNA3*, *FGF5*, *EFNA4*, *IGF1*, *EFNA2*, *FGF9*, *FGF13*, *FGF17*, *PDGFB*, *NGF*, *ANGPT4*, *FGF7*, *FGF22*, *FGF16*, *FGF23*, *FGF19*, *FGF20*, *FGF8* and *VEGFD*). B) in the receptor node of the circuits within *Focal adhesion pathway* the *VEGFA* potentially shares the signal transduction role with other 12 proteins (*PDGFA*, *HGF*, *VEGFB*, *PDGFC*, *EGF*, *PDGFD*, *PGF*, *VEGFC*, *IGF1*, *PDGFB* and *VEGFD*). C) in the 6 signaling circuits belonging to the *HIF signaling pathway* and *VEGF signaling pathway* the protein *VEGFA* is the only signal transducer in the node.

The same modeling strategy used with bevacizumab can be applied to simulate the effect of other drugs. Recently, a drug repurposing *in silico* experiment which combines human genomic data with mouse phenotypes has suggested the possible utility of a number of drugs with different indications (see Table 3) for potential glioblastoma treatment (60). The intensity and the degree of heterogeneity in the response are very variable across the 8 drugs tested here. At the scale we tested the drugs, there are no correlations neither between the number of genes targeted by the drug and the intensity of the effect nor between the number of circuits potentially affected and the intensity of the effect. For example, Pentoxifylline targets 4 genes (*PDE4B, ADORA1, PDE4A* and *ADORA2A*) which participate in a total of 111 circuits and the fold change caused in the circuit activity after simulating its effect is comparatively low (log fold change < 0.05 for all the cell types; see Supplementary Figure 1), while the simulation of the effect of Fenofibrate, which targets only one gene *PPARA* that participates in only 32 circuits renders a comparatively high effect (log fold change > 2 for all the cell types; see Supplementary Figure 1). It is interesting to note that, depending on the case, the different drug effects simulated can affect to a larger or to a smaller number of cells with distinct intensity in their impacts on the activity of the signaling circuits affected, but always, no matter what drug is simulated, there are some cells that manage to escape to the inhibitory effect of the drug.

**Table 3.**
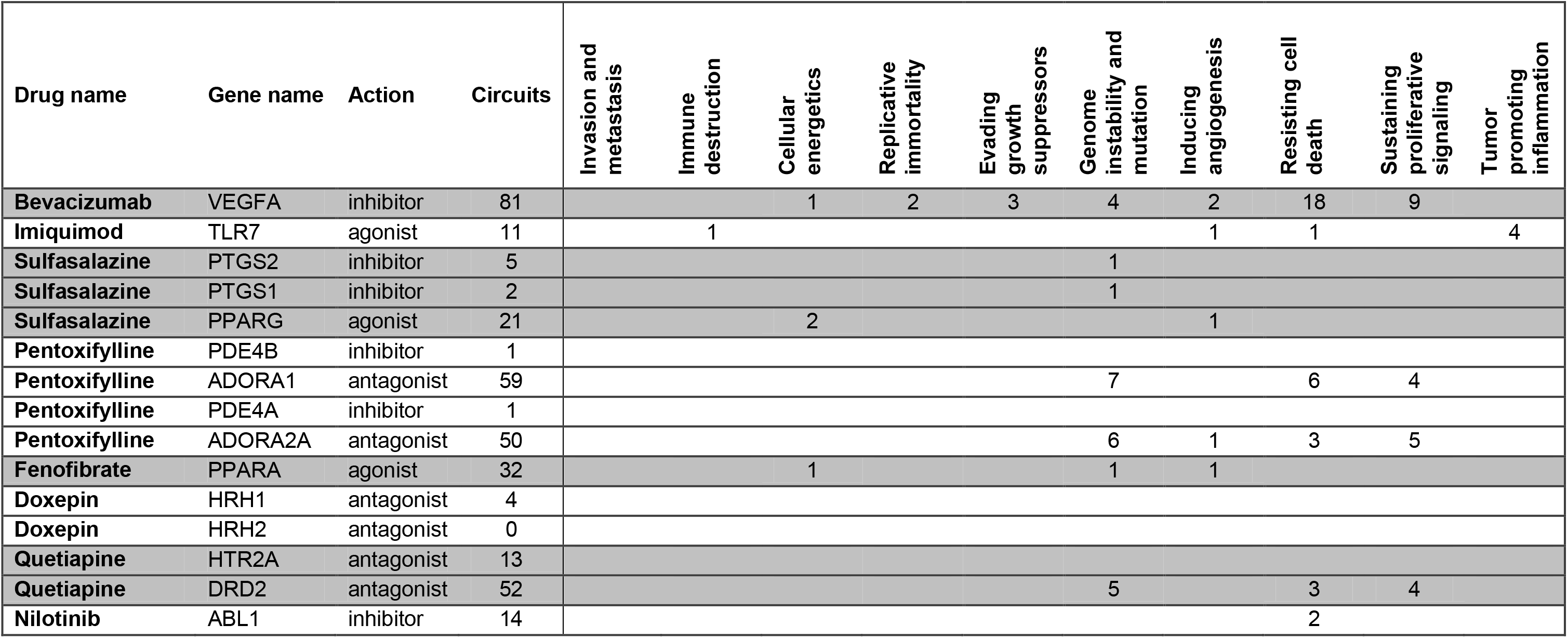
The eight drugs whose effect on the neoplastic population cell has been simulated. The hallmarks affected are displayed in the columns of the right of the table.

## Discussion

The goal of most scRNA-seq publications revolves around the characterization of cell populations, which can be accurately achieved using only a subset of the total number of genes (those displaying the highest variability across cells). However, the use of mechanistic models to estimate global signaling circuit activity profiles for individual cells requires of reasonably accurate measures of the expression levels of all the genes involved in the signaling circuits. Dropout events, quite common in scRNA-seq experiments (46), are particularly problematic given that taking by mistake a missing value by a real zero value can cause erroneous determinations of the inferred activity of the circuits. Thus, we explored the performance of three different imputation methods in producing cell-specific profiles of signaling circuit activity whose clustering resulted in a grouping similar to that observed and validated in the original glioblastoma study. Here, two machine learning based methods, DRImpute and MAGIC, produced a clustering compatible with the original validated clustering, and specifically, Drimpute, the method of choice, rendered clusters with a similar shape as well (see Figure 1).

Focusing on neoplastic cells, the existence of three different clusters is also apparent at the level of functional profiles, which suggests the existence of different functional behaviors. Several attempts to stratify glioblastoma patients have been proposed by discriminating different subtypes according to different properties, such as patient survival (61) mutational status of some genes (62) or the tumor microenvironment (63). In the most used classification, glioblastoma tumors were divided into three subtypes (from less to more aggressive: classical, proneural, and mesenchymal) based on the signature of 50 genes (55). Although this conventional subtyping provides an approximate descriptor of tumor aggressiveness, subtyping based on functional profiles related to cancer hallmarks provides an interesting alternative for the stratification of glioblastoma that offers, in addition, a mechanistic description on the functional activity of the tumor. Actually, it has been reported in neuroblastoma that signaling pathway models used as biomarkers outperform traditional biomarkers as predictors of patient survival (26).

Among pathways that are commonly altered in all three clusters we find well known factors contributing to carcinogenesis, such as those related to hypoxia (HIF-1, SOD2), cancer stem cells (CSCs), cell cycle proteins, like CDK family, signal transduction pathways and hormone signaling (64–66). Moreover, these are mainly related to *Sustaining proliferative signaling, Enabling replicative immortality* and *Resisting cell death* hallmarks that can be defined as the core cell functions involved in glioblastoma initiation and proliferation.

Each cluster exhibits a characteristic deregulation of pathways, however, cluster 2 barely has four unique sub-pathways, all related to common hallmarks *Resisting cell death* and *Sustaining proliferative signaling*. Neoplastic clusters 1 and 3, but no cluster 2, exhibit *Genome instability*, a hallmark observed in almost all sporadic human cancers, including glioblastoma (67,68). Besides, cluster 1 and 3, show deregulated *Cellular energetics*, a process that has been suggested as a suitable target for tumor cell elimination (69). Furthermore, both clusters show pathways associated with matrix metalloproteins and Snail family that have been linked to cancer invasion and metastasis also in glioblastoma (70–72). Interestingly, only cluster 3 may be avoiding immune destruction due to the deregulation of *Toll-like receptor signaling pathway*. Glioblastoma is known to have a strongly immunosuppressive microenvironment, thus blocking these cells by activating downstream TLR signaling pathways can reduce tumor growth and disrupt CSC self-renewal (73,74).

We have demonstrated that not all the cells in a tumor are driven by the same cancer processes, and that those alterations can define subpopulations that may confer tumors different aggressiveness and invasion abilities, highlighting the relevance of heterogeneity, beyond the widely accepted stratification of glioblastoma in three/four subtypes (75,76).

The emergence of mechanisms of resistance in targeted therapies has been attributed to either the selection of rare pre-existing genetic alterations upon drug treatment (77) or the transient acquisition of a drug-refractory phenotype by a small proportion of cancer cells through epigenetic modifications (78). In both cases, these alterations would be detectable in the expression of the corresponding genes. An interesting property of mechanistic models is that they can be used to model the effect of an intervention over the system studied (25,35). Thus, the use of mechanistic models on single cell transcriptomic data offers for the first time the possibility of modeling the effects of a targeted drug in individual cells. From Figure 4 it becomes apparent that low-responder cells have a constitutive level of *VEGFA* lower than highly-responder cells, either with the relevant role of signal transduction taken by other proteins to keep the circuit active or simply with the circuit already inactive. Thus, the mechanistic model provides a simple explanation of the molecular mechanisms behind the differential effect of drugs over cells with different signaling profiles that ultimately cause different functional strategies.

Supplementary Figure 1 depicts the simulated response of individual cells to the treatment with different targeted drugs. Despite the variety of effects that cause in the cells, It is worth noticing that there is always a group of cells that manage to escape to the inhibition of the drug. The heterogeneity observed in the cell population in terms of use of different strategies to activate the essential cancer hallmark through different signaling circuits, produces a consequent diversity in the response to drugs. Although the number of drugs simulated is relatively low, given that drug repurposing was beyond the scope of this paper, the results obtained here suggest that the escape of a relatively small number of cells to the effect of the drug could be a relatively frequent event that occurs as a natural consequence of the heterogeneity in the signaling strategies followed by the cell population. When this subpopulation becomes dominant some time later, it results to be resistant to the drug.

## Conclusions

The use of mechanistic models provides a detailed insight into the functional strategies used by tumors to proliferate and open new avenues for the design of interventions *a la carte*. The extension of this analytic approach to single cell transcriptomic data allows an unprecedented detail on how cancer cells display different functional strategies to proliferate that have consequences in their respective vulnerabilities to targeted therapies.

Although the existence of resistant clones in a tumoral cell population is well known, the specific mechanisms used by resistant cells to escape from the inhibitory effects of targeted therapies remain unknown yet. Mechanistic models offer for the first time a plausible and contrastable hypothesis on how and why some cells are insensitive to treatments, illustrated here with bevacizumab in the case of glioblastoma. The use of this modeling strategy offers a systematic way for detecting tumoral cells that may be resistant to specific targeted treatments. Conversely, the same models could be used to find an alternative treatment for resistant drugs. In fact our results suggests that the search for new, more efficient therapeutic targets would be very benefited of the use of mechanistic models that guide to the intervention points with more likelihood of success in inhibiting the proliferation of the largest possible part of the spectrum of functional strategies in the tumor cell ecosystem.

## Material and methods

### Data and primary processing

A large scRNA-seq dataset containing 3589 cells of different types obtained in four patients from a glioblastoma study (45) was downloaded from GEO (GSE84465). Cells corresponded to the tumor, and to the periphery of the tumor.

Count values for the scRNA-seq were downloaded from GEO and subjected to the three imputation methods, MAGIC (48), SCimpute (49) and Drimpute (50), as implemented in the corresponding software packages. Each method has its own preprocessing pipeline explained in the corresponding publication.

Once imputed, samples were log-transformed and a truncation by quantile 0.99 was applied. Finally, the values were normalized between 0 and 1, as required by the downstream functional analysis with *Hipathia*.

### Hipathia mechanistic model

The *Hipathia* method uses KEGG pathways (79) to define circuits that connect any possible receptor protein to specific effector proteins. Gene expression values are used in the context of these circuits to model signaling activity, that ultimately trigger specific cell activities, are as described in (24). A total of 98 KEGG pathways involving a total of 3057 genes that form part of 4726 nodes were used to define a total of 1287 signaling circuits. The intensity value of signal transduced to the effector is estimated by the following recursive formula:

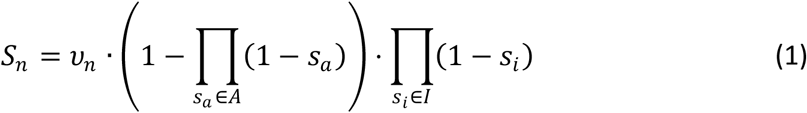

where *S_n_* is the signal intensity for the current node *n*, *v_n_* is its normalized gene expression value, *A* is the set of activation signals (*s_a_*), arriving to the current node from activation edges, *I* is the set of inhibitory signals (*s_i_*) arriving to the node from inhibition edges (24).

The *Hipathia* algorithm (27) is implemented as an R package available in Bioconductor (52) as well as in a web server (80) and as a Cytoscape application (81).

### Differential signaling activity

Two groups of circuit activity profiles can be compared and the differences in activity of any circuit can be tested by means of different tests, among which *limma* (82) has demonstrated to be very efficient.

### Signaling circuits associated to cancer hallmarks

Each effector is known to be associated to one or several cell functions. This information is extracted both from the Uniprot (83) and Gene Ontology (84) annotations corresponding to the effector gene (24). However, in some cases the annotations are too ambiguous or make reference to roles of the gene in many different conditions, tissues, developmental stages, etc., making thus difficult to understand its ultimate functional role. In addition, in this study the activity of signaling circuits relevant in cancer is particularly interesting. Since a number of these effector genes has been related specifically with one or several cancer hallmarks (34) in the scientific literature, the CHAT tool (54), a text mining based application to organize and evaluate scientific literature on cancer that allows linking gene names with cancer hallmarks.

### Subtyping of cancer cells

The *SubtypeME* tool from the GlioVis data portal (56) was used to obtain the subtype of cancer (classical, proneural or mesenchymal), based on the signature of 50 genes (55). This tool provides three methods to assign subtype: Single sample Gene Set Enrichment Analysis (ssGSEA), K-nearest neighbors (k-NN) and Supported vector machine (SVM). Subtype was assigned when at least two of the methods made an identical subtype prediction.

## Acknowledgements

This work is supported by grants SAF2017-88908-R from the Spanish Ministry of Economy and Competitiveness and “Plataforma de RecursosBiomoleculares y Bioinformáticos” PT17/0009/0006 from the ISCIII, both co-funded with European Regional Development Funds (ERDF) as well as H2020 Programme of the European Union grants Marie Curie Innovative Training Network "Machine Learning Frontiers in Precision Medicine” (MLFPM) (GA 813533) and “ELIXIR-EXCELERATE fast-track ELIXIR implementation and drive early user exploitation across the life sciences” (GA 676559). The results shown here are in part based upon data generated by the TCGA Research Network: https://www.cancer.gov/tcga.

