## Supplementary Figure 1 for "Mechanistic models of signaling pathways deconvolute the functional landscape of glioblastoma at single cell resolution"

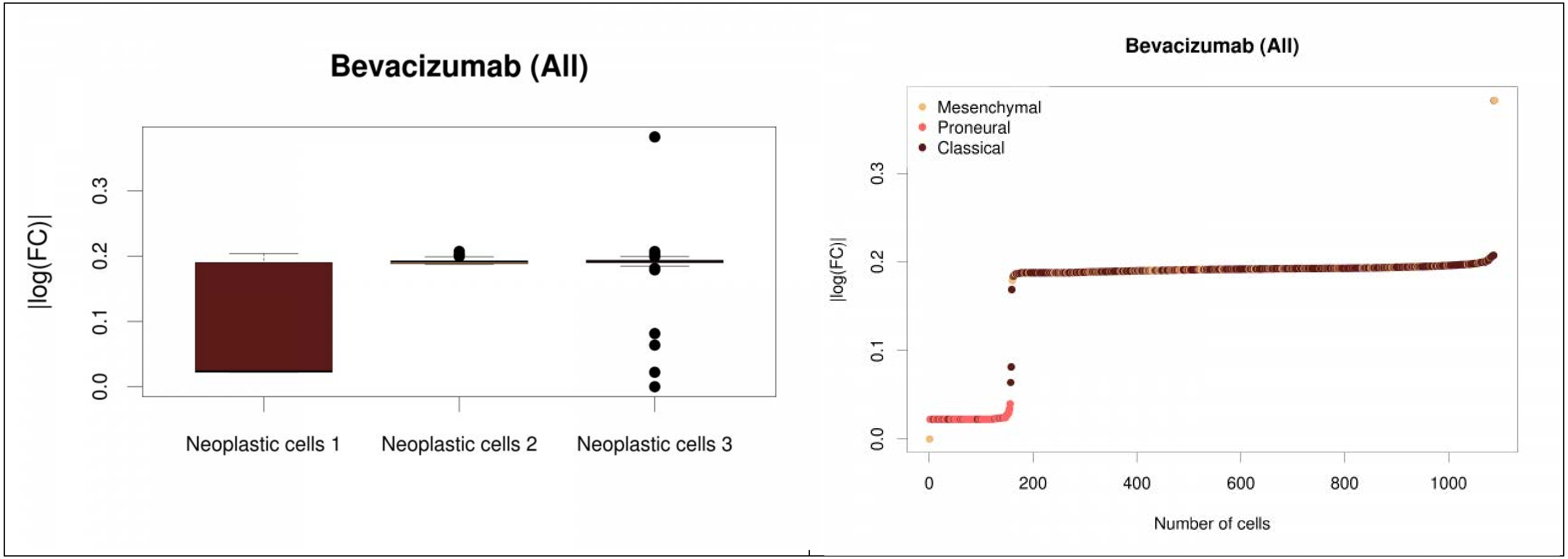

Bevacizumab (All)
|log(FC)|
0.0
0.1
0.2
0.3
Neoplastic cells 1
Neoplastic cells 2
Neoplastic cells 3
Bevacizumab (All)
Mesenchymal
Proneural
Classical
|log(FC)|
0.0
0.1
0.2
0.3
0
200
400
600
800
1000
Number of cells

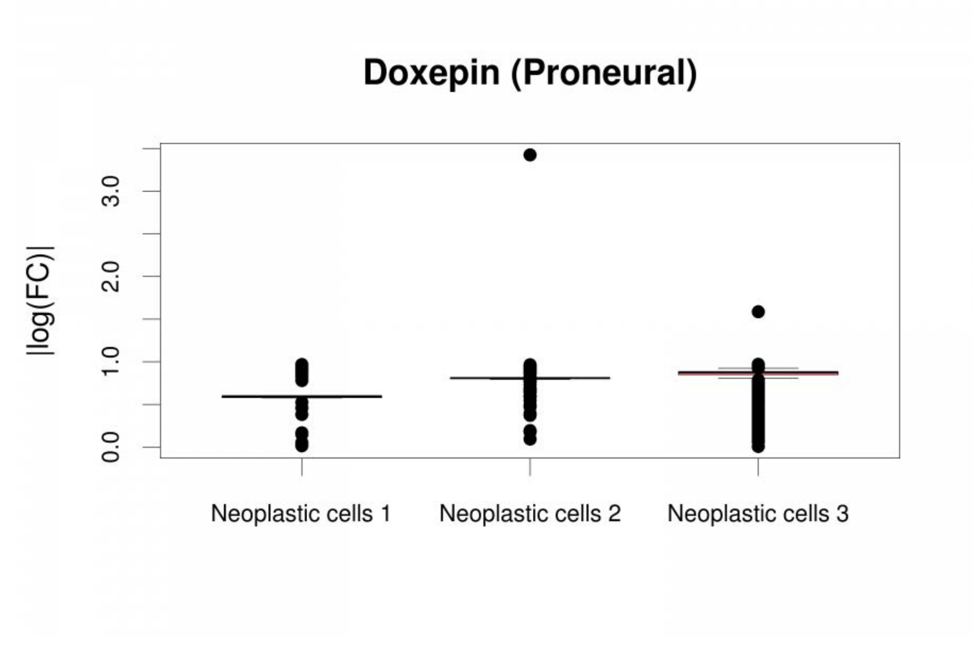
Doxepin (Proneural)
|log(FC)|
0.0
1.0
2.0
3.0
Neoplastic cells 1
Neoplastic cells 2
Neoplastic cells 3

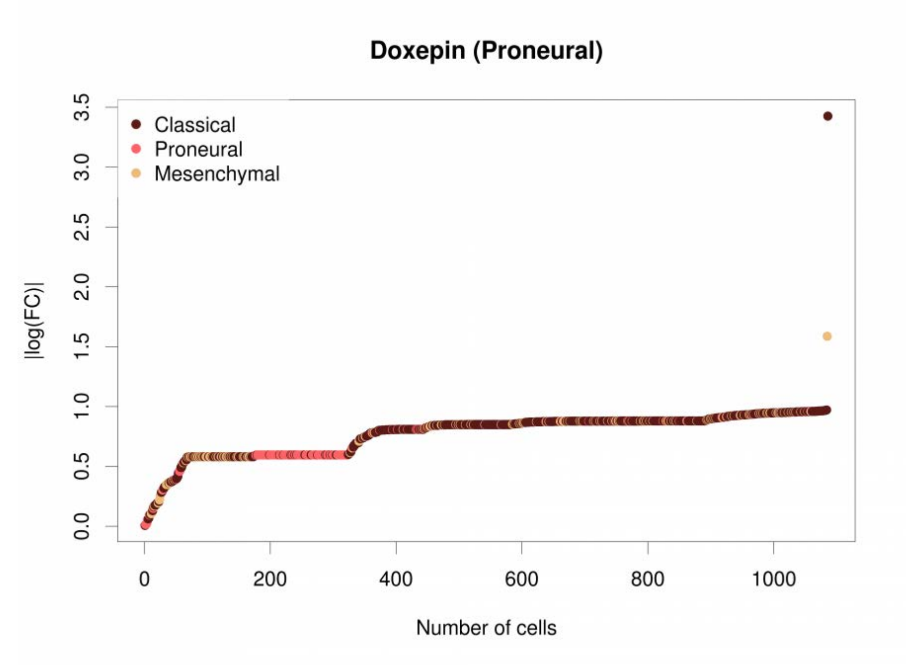
Doxepin (Proneural)
Classical
Proneural
Mesenchymal
|log(FC)|
0.0
0.5
1.0
1.5
2.0
2.5
3.0
3.5
0
200
400
600
800
1000
Number of cells

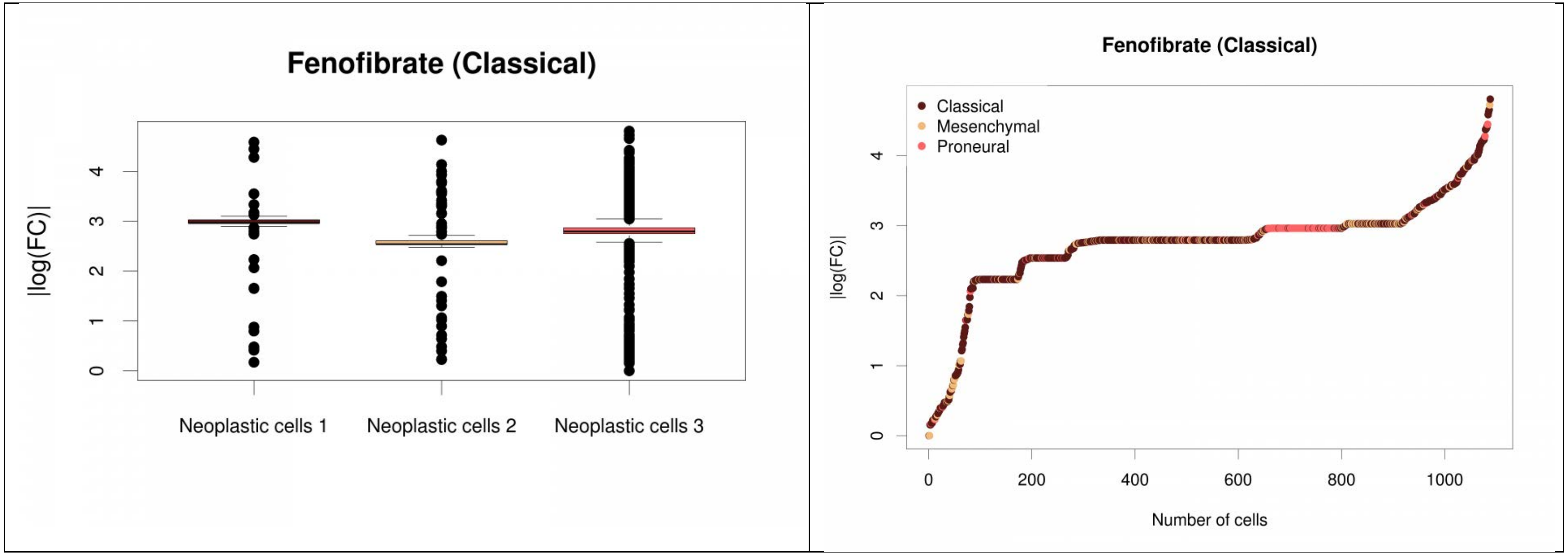

Fenofibrate (Classical)
|log(FC)|
0
1
2
3
4
Neoplastic cells 1
Neoplastic cells 2
Neoplastic cells 3
Fenofibrate (Classical)
Classical
Mesenchymal
Proneural
|log(FC)|
0
1
2
3
4
0
200
400
600
800
1000
Number of cells

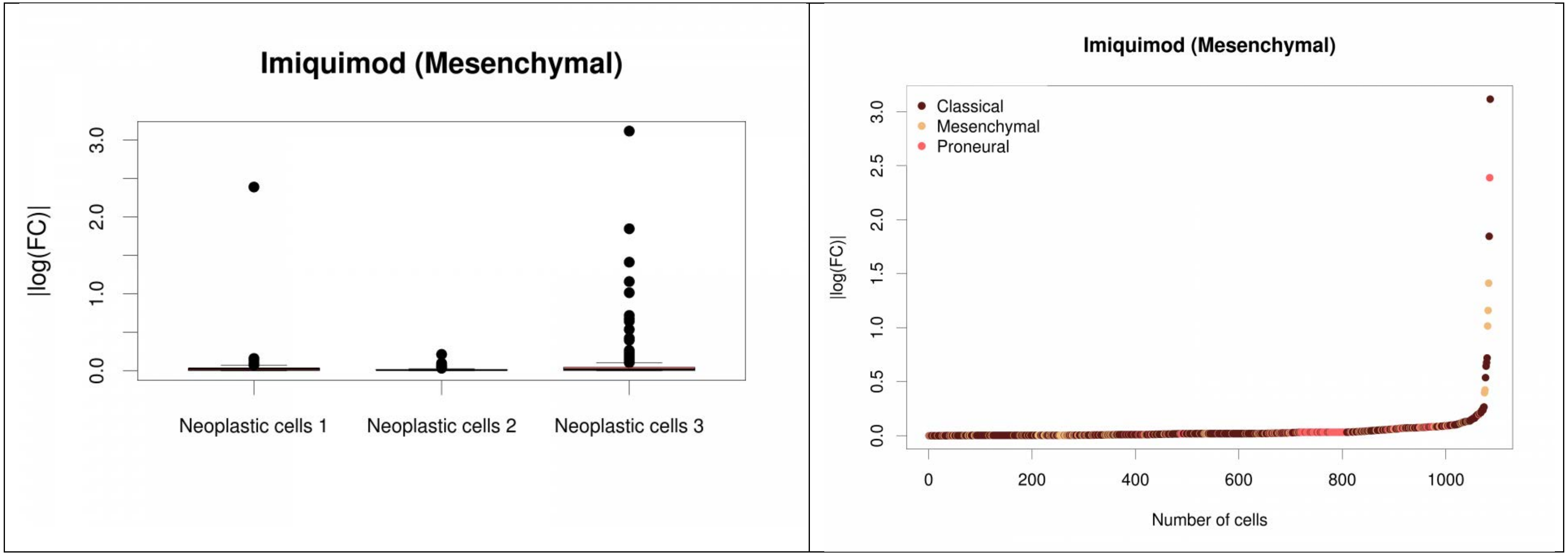

Imiquimod (Mesenchymal)
|log(FC)|
0.0
1.0
2.0
3.0
Neoplastic cells 1
Neoplastic cells 2
Neoplastic cells 3
Imiquimod (Mesenchymal)
Classical
Mesenchymal
Proneural
|log(FC)|
0.0
0.5
1.0
1.5
2.0
2.5
3.0
0
200
400
600
800
1000
Number of cells

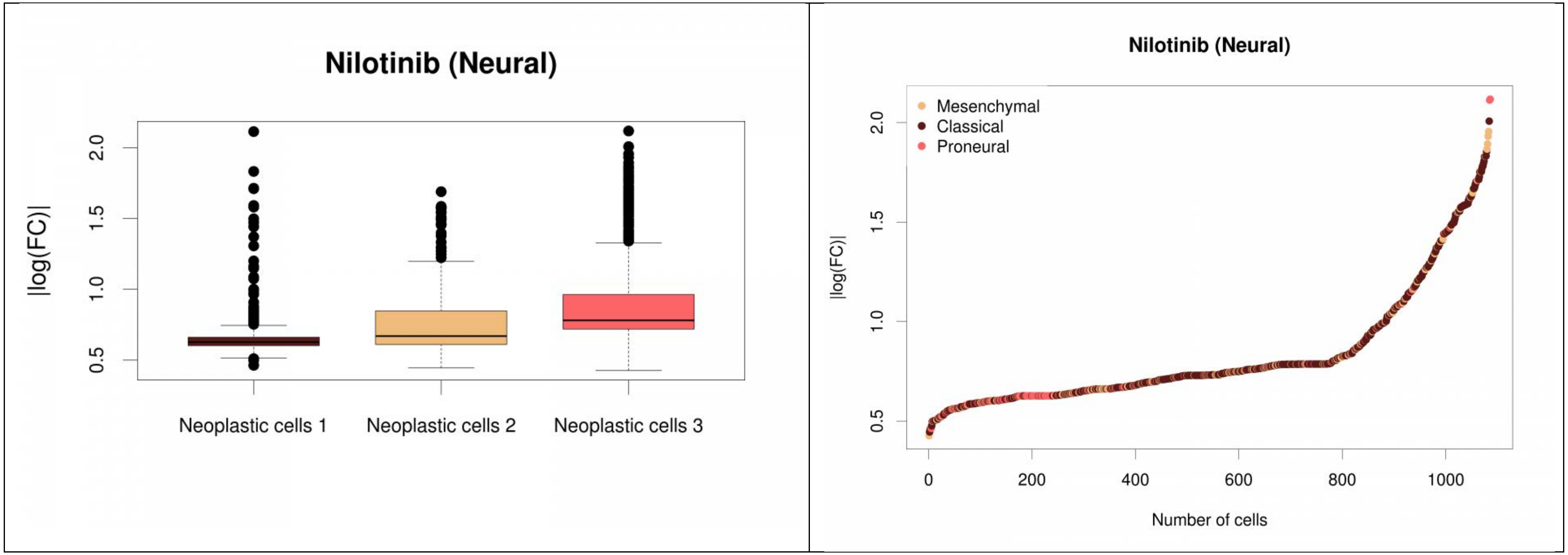
Nilotinib (Neural)
|log(FC)|
2.0
1.5
1.0
0.5
Neoplastic cells 1
Neoplastic cells 2
Neoplastic cells 3
Nilotinib (Neural)
Mesenchymal
Classical
Proneural
|log(FC)|
0
200
400
600
800
1000
Number of cells

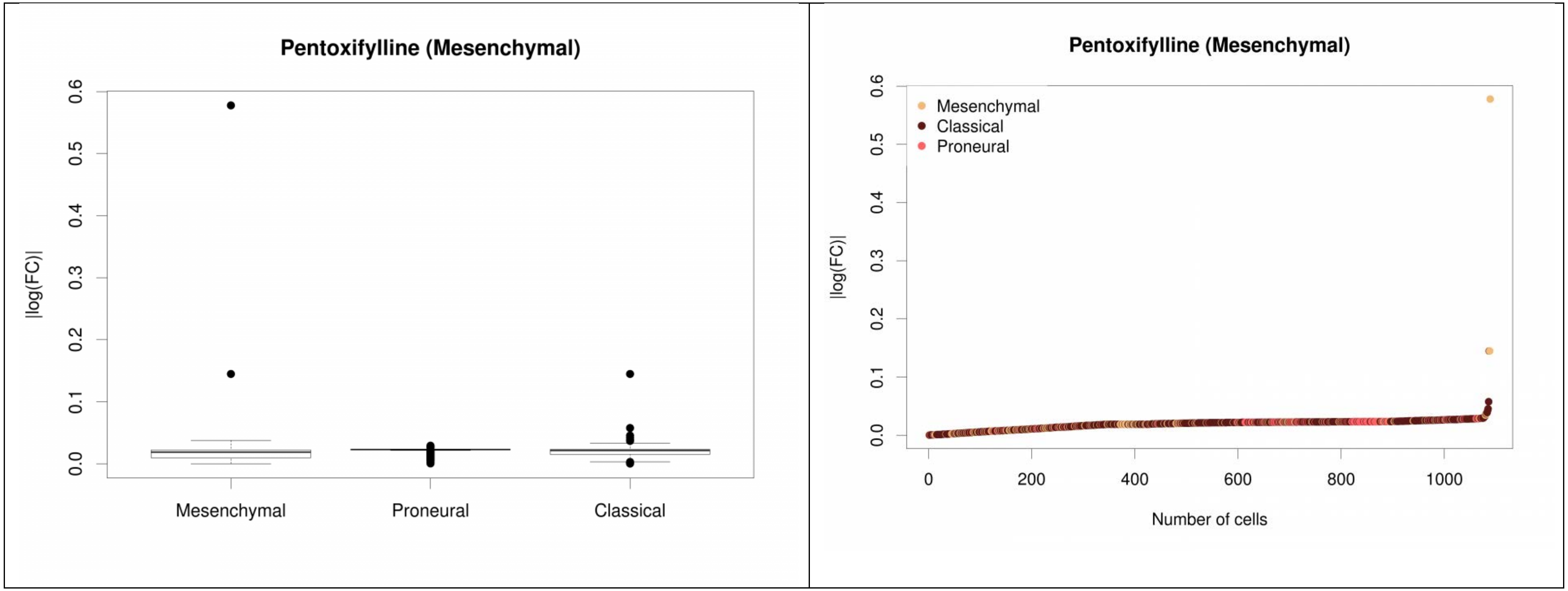
Pentoxifylline (Mesenchymal)
|log(FC)|
0.0
0.1
0.2
0.3
0.4
0.5
0.6
Mesenchymal
Proneural
Classical
Pentoxifylline (Mesenchymal)
|log(FC)|
Mesenchymal
Classical
Proneural
0
200
400
600
800
1000
Number of cells

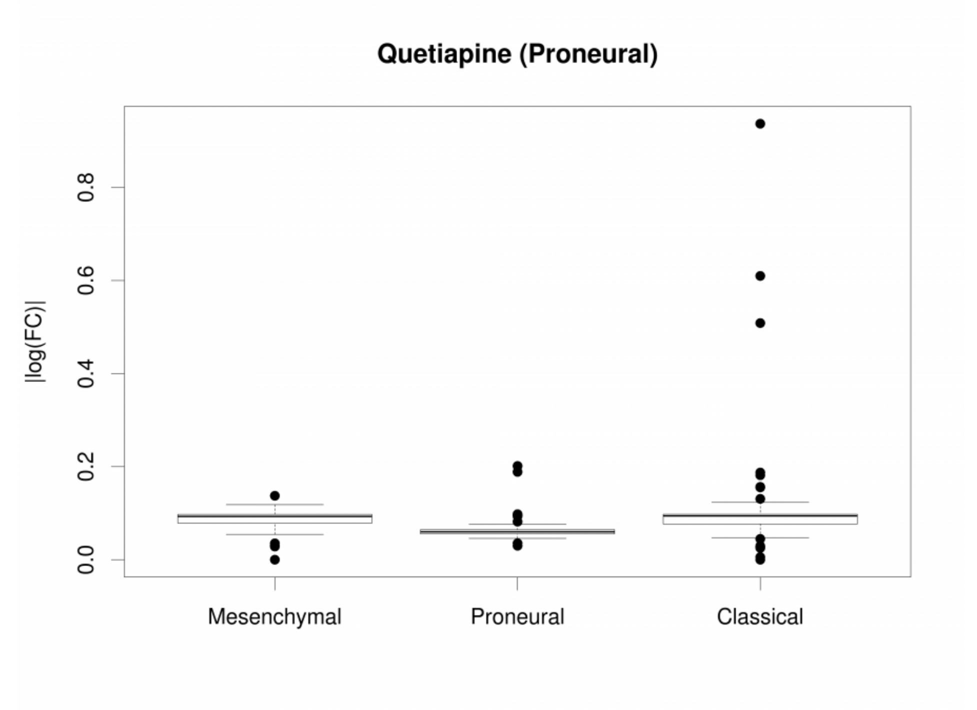

Quetiapine (Proneural)
|log(FC)|
0.0
0.2
0.4
0.6
0.8
Mesenchymal
Proneural
Classical


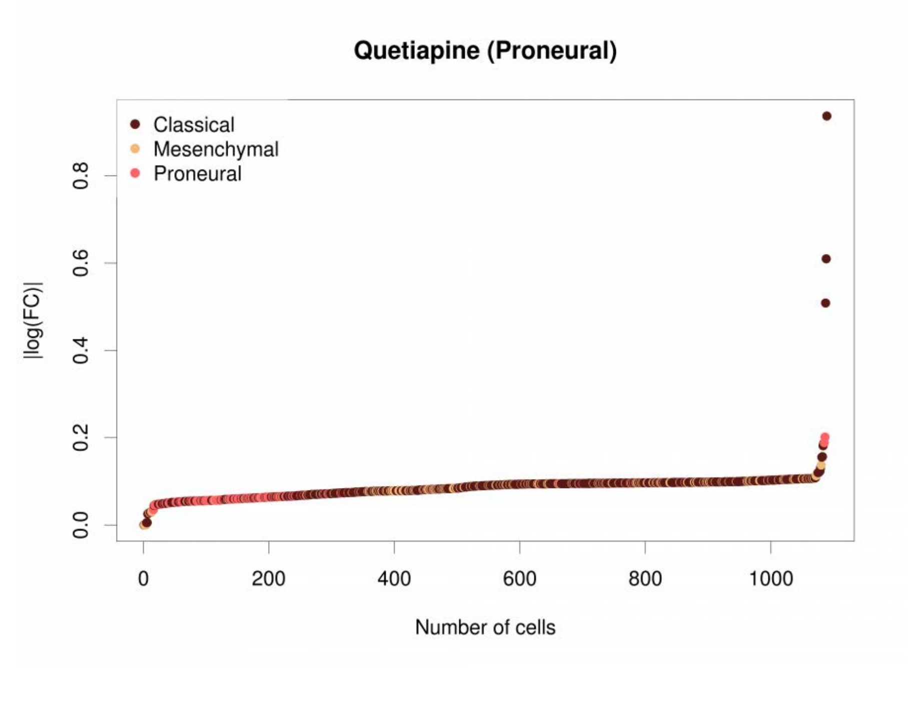

Quetiapine (Proneural)
Classical
Mesenchymal
Proneural
|log(FC)|
0.0
0.2
0.4
0.6
0.8
0
200
400
600
800
1000
Number of cells

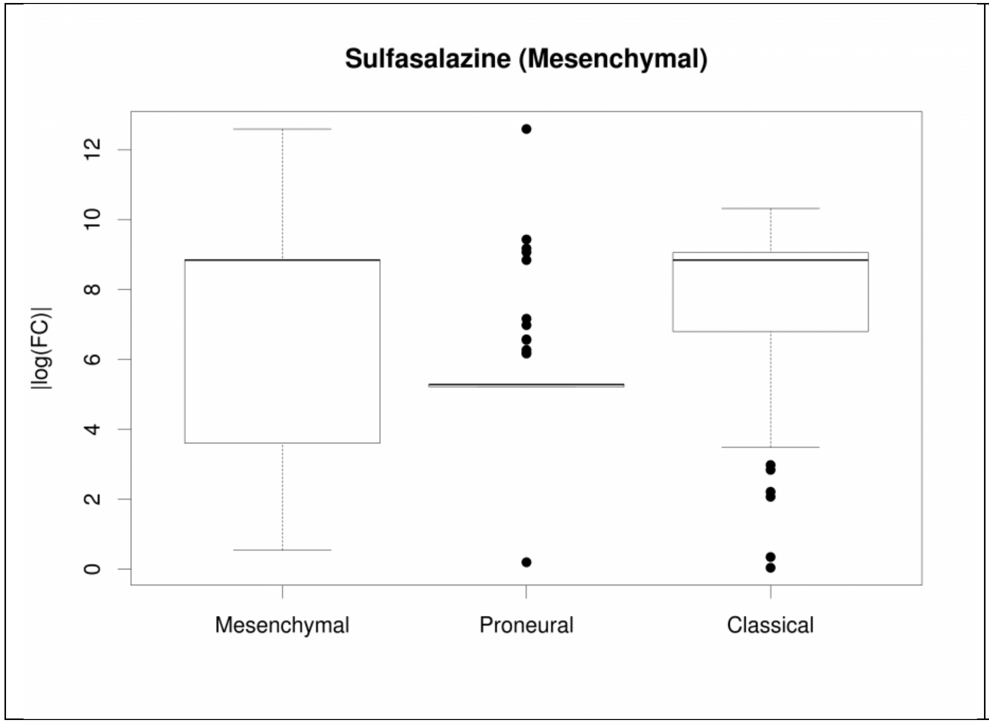

Sulfasalazine (Mesenchymal)
|log(FC)|
0
2
4
6
8
10
12
Mesenchymal
Proneural
Classical


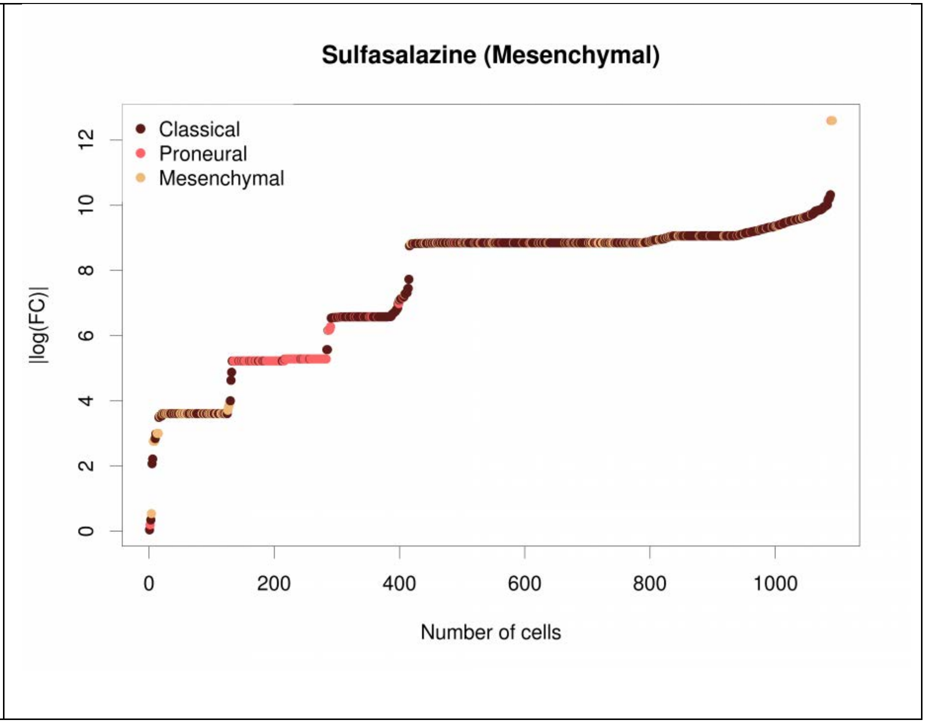

Sulfasalazine (Mesenchymal)
Classical
Proneural
Mesenchymal
|log(FC)|
0
2
4
6
8
10
12
0
200
400
600
800
1000
Number of cells
